# A minimal “push-pull” bistability model explains oscillations between quiescent and proliferative cell states

**DOI:** 10.1101/239897

**Authors:** Sandeep Krishna, Sunil Laxman

## Abstract

A minimal model for oscillating between quiescent and growth/proliferation states, dependent on the availability of a central metabolic resource, is presented. From the yeast metabolic cycles (YMCs), metabolic oscillations in oxygen consumption are represented as transitions between quiescent and growth states. We consider metabolic resource availability, growth rates, and switching rates (between states) to model a relaxation oscillator explaining transitions between these states. This frustrated bistability model reveals a required communication between the metabolic resource that determines oscillations, and the quiescent and growth state cells. Cells in each state reflect memory, or hysteresis of their current state, and “push-pull” cells from the other state. Finally, a parsimonious argument is made for a specific central metabolite as the controller of switching between quiescence and growth states. We discuss how an oscillator built around the availability of such a metabolic resource is sufficient to generally regulate oscillations between growth and quiescence, through committed transitions.

## Introduction

While all cells can exist in a variety of states, two opposite ends of the spectrum are the “growth” state (leading to mitotic division and proliferation), and a non-proliferative “quiescent” state. The quiescent state, operationally defined here as a reversibly non-dividing state, is the predominant state of all living cells (Lewis and Gattie, 1991; Gray *et al.*, 2004). Understanding how cells reversibly transition from a quiescent state, to a growth state coupled with cell division and proliferation (henceforth called “growth” in this manuscript) is therefore a fundamental biological question. Current explanations for how cells commit to growth and cell division account for metabolic regulation, biomolecule synthesis, and regulated progression through the cell cycle, presenting multiple, integrated mechanisms of information transfer within a cell that lead to the eventual growth outcome.

However, when a population of genetically identical cells are present in a uniform environment, how can individual cells within such a population decide to switch between a quiescent (effective “G0”) state and a growth/proliferation state? Indeed, such heterogeneity of cell states within populations is widely observed and acknowledged. Numerous examples exist in nearly all systems studied, from simple eukaryotes like the budding yeast, to complex mammalian systems (Cooper, 1998, 2003; Coller *et al.*, 2006; Daignan-Fornier and Sagot, 2011; Klosinska *et al.*, 2011; De Virgilio, 2012; Dhawan and Laxman, 2015), with multiple molecular events correlating with transitions between growth and quiescence. For any population transitioning into either of these states, experimentalists have asked: (i) what hallmarks allow discrimination between actively proliferating and G0 cells? (ii) how do cells transit back and forth between these two states? And (iii) how are different signals processed and integrated into an appropriate cellular response? The regulation of the final cellular outcome occurs at multiple levels, including differential gene expression programs, and signaling responses to growth factors, which can be different depending upon the type of cell or organism studied. At its very core, however, this transition between quiescent and growth states is a metabolic problem; cells must be in a metabolic state capable of committing to growth/proliferation, and must sense this state, which the pushes cells towards growth. Indeed, several lines of evidence now reiterate a primary metabolic determinant for cells committing to a growth state (exiting quiescence), or remaining in a quiescent state (Futcher, 2006; Daignan-Fornier and Sagot, 2011; Laporte *et al.*, 2011; Cai and Tu, 2012; De Virgilio, 2012; Lee and Finkel, 2013; Dhawan and Laxman, 2015; Kalucka *et al.*, 2015; Kaplon *et al.*, 2015). While multiple factors can regulate the transition between quiescence and growth, all such studies suggest that without this core metabolic transformation, switching states is impossible. Given this absolute metabolic requirement to switch to growth, if there is an isogenic (“identical”) population of cells present in a uniform environment, how can there be a two-state outcome where some cells undergo growth/proliferation, while the rest remain quiescent?

Surprisingly, there are few rigorous theoretical, mathematical models that attempt to provide a conceptual framework sufficient to explain this, and suggest experimentally testable predictions. This is in contrast to the extensive, elegant, and often prescient models that have been built to explain progress through the classical cell division cycle (CDC), by incorporating existing experimental data of phase specific cell-cycle activators and inhibitors (Tyson and Novak, 2001; Tyson *et al.*, 2003; Ferrell *et al.*, 2009; Tyson and Novák, 2015). Such modeling of the CDC has a long history (examples include (Goldbeter, 1991; J, 1991; Norel and Agur, 1991; Novák and Tyson, 1993; Ferrell *et al.*, 2009; Tyson and Novak, 2015)), and these types of theoretical studies have revealed biological possibilities that were experimentally determined only much later (such as (Cross *et al.*, 2002; Pomerening *et al.*, 2003; Wei *et al.*, 2003; Mirchenko and Uhlmann, 2010)). Given this, there is considerable value in building coarse-grained but rigorous theoretical models to understand switching between quiescence and growth states. In such a model, the switching between quiescence and growth states could be treated as a biological oscillation (Tyson *et al.*, 2003; Novák and Tyson, 2008; Tsai *et al.*, 2008; Ferrell *et al.*, 2009), while considering a dependence on a metabolic “resource” as a driver of the oscillator. For building such a model, we therefore require extensive experimental data from biological systems where metabolic oscillations are demonstrably closely coupled with exiting quiescence/entering the CDC.

Such data are readily available from the budding yeast, S. cerevisiae. Yeast have been the instrumental cellular model in revealing processes that define both the CDC (Hartwell, 1974), and the quiescence cycle (Gray *et al.*, 2004; Daignan-Fornier and Sagot, 2011; Daignan-Fornier B and Sagot I, 2011; De Virgilio, 2012; Dhawan and Laxman, 2015). The classical CDC involves progression through the G1, S, and G2/M phases. In contrast, during a quiescence (or effective “G0”) cycle, cells remain non-dividing, but can exit quiescence and enter the G1 phase of the cell cycle to subsequently complete the CDC.

Experimentally dissecting specific processes driving entry into, and exit from, quiescence (into the CDC) is challenging in asynchronous, heterogeneous cultures of cells. However, synchronized yeast populations in well-mixed cultures (as manifest by oscillations in oxygen consumption) have long been observed and studied using batch and chemostat conditions limited for a carbon source (glucose or ethanol), which are subsequently fed continuously with limited concentrations of glucose or ethanol (Chance *et al.*, 1964; Hommes, 1964; Hess and Boiteux, 1971; Satroutdinov *et al.*, 1992; Keulers *et al.*, 1996; Jules *et al.*, 2005; Lloyd and Murray, 2005). Gene expression studies from such glucose-limited yeast metabolic cycles or oscillations (we will utilize the term YMC henceforth in this manuscript for consistency) showed that a majority of the genome is expressed highly periodically, further revealing a molecular organization of growth and quiescent states (Klevecz *et al.*, 2004; Tu *et al.*, 2005; Futcher, 2006; Mellor, 2016). In general, both the shorter (Klevecz *et al.*, 2004; Murray *et al.*, 2007), and the longer oxygen consumption oscillations in yeast (Tu *et al.*, 2005) showed this general pattern. Notably, genes associated with biosynthesis and growth (comprehensively further described in (Brauer *et al.*, 2008)) typically peak during a high oxygen consumption phase in the YMC (Tu *et al.*, 2005; Rowicka *et al.*, 2007; Slavov and Botstein, 2011, 2013), while genes that mark autophagy, vacuolar function and a “quiescence” state peak during a steady, low oxygen consumption phase. Strikingly, in these continuous YMC cultures, cell division is tightly gated to a temporal window. Cells divide synchronously only once during each metabolic cycle (Küenzi and Fiechter, 1969; Tu *et al.*, 2005; Robertson *et al.*, 2008; Laxman *et al.*, 2010) and remain in a non-dividing state during the rest of the cycle. The non-dividing population in the low oxygen consumption phase exhibits typical hallmarks of quiescent cells (Tu *et al.*, 2005, 2007; Shi *et al.*, 2010; Cai *et al.*, 2011; Shi and Tu, 2013; Dhawan and Laxman, 2015). Furthermore, in each YMC, during the tight temporal window when cells do divide, the culture has two, visibly distinct sub-populations: dividing and nondividing (Tu *et al.*, 2005; Robertson *et al.*, 2008; Laxman *et al.*, 2010). These data have suggested a close coupling between the metabolic and the cell division cycles. Importantly, the YMC itself is metabolite/nutrient regulated, and controlled by the amount of available glucose. The distinct phases of the YMC correspondingly show a separation of metabolic processes (Tu *et al.*, 2005, 2007; Murray *et al.*, 2007; Machné and Murray, 2012), and several lines of evidence suggest that key metabolite amounts are critical for entering or exiting a proliferative or a non-proliferative state (Murray *et al.*, 2003, 2007; Tu *et al.*, 2007; Shi *et al.*, 2010; Cai *et al.*, 2011; Machné and Murray, 2012; Mellor, 2016). These studies collectively indicate the following: (i) a separation of two states (proliferative, and effectively G0) in cell populations, dependent on metabolic states, and (ii) a loose metabolic framework within which it may be possible to study transitions between quiescence and growth transitions. Thus, these studies provide extensive experimental data using which a theoretical, mathematical model can be built to sufficiently explain oscillations between a “quiescent” state and a “growth” state.

Here, we use existing data from these YMCs to build a robust, general model for oscillations between a quiescent and a growth state. Importantly, the model necessitates the requirement of a tripartite communication - between the metabolic resource, the quiescent cells, and the cells exiting quiescence and entering growth - in order for the cells to sustain oscillation between these two states. The model oscillations depend on an underlying bistability, suggesting that cells in either state exhibit hysteresis, or memory, of their states. Finally, using this model, we show how two central metabolites, thought to be critical for entry into a growth state, satisfy the required criteria for the currency that controls oscillations between these two cell states. Collectively, we provide a coarse-grained, sufficiency model to explain general principles of how cells can oscillate between a quiescent and growth state, depending upon amounts and utilization of an internal metabolic currency.

## Results

### Apparent bistable states during yeast metabolic cycles

Yeast cells grown to a high cell density (in batch culture mode) in a chemostat, and when subsequently fed limited amounts of glucose medium, spontaneously undergo robust oscillations in oxygen consumption (YMCs) (Figure 1A) and (Klevecz *et al.*, 2004; Tu *et al.*, 2005; Murray *et al.*, 2007; Silverman *et al.*, 2010; Burnetti *et al.*, 2015), with the period of each oscillation ranging from ~2.5-5 hours (Figure 1A). For these oscillations to occur, the batch culture typically needs to first be starved for a few hours (Figure 1A), during which time all glucose is depleted, and all cells enter a non-dividing state (although the extended starvation is not an absolute requirement, as observed historically in breweries). After starvation, when cells are continuously provided limited glucose in the medium, the oscillations in oxygen consumption spontaneously start and continue indefinitely (Figure 1A). Comprehensive gene expression analysis across these longer-period oscillations (1.5-4.5 hr cycles) has revealed highly periodic transcript expression (Tu *et al.*, 2005; Rowicka *et al.*, 2007), and proteins encoded by these transcripts can be binned into three general classes (Figure 1B, 1C). These represent “growth genes” during the high oxygen consumption phase, followed by the rapid decrease in oxygen consumption coupled with “cell division” (Figure 1B, 1C) (Kudlicki *et al.*, 2007; Rowicka *et al.*, 2007). The cells exhibiting the “growth” signature during the high oxygen consumption phase all go on to enter and complete the CDC (42). Finally, the YMC enters a state of ~stable oxygen consumption, where the gene expression profile revealed a “quiescent”-like state (Figure 1B, 1C). Mitotic cell division is tightly gated only to a narrow window (Figure 1B, 1C). Interestingly, in this phase, only a fixed fraction of the cells (~35%) (and not all cells) divide during each cycle (Figure 1D). During the stable oxygen consumption phase, there are almost no budding cells observed (Figure 1D). Note: given that this is a controlled chemostat system, the overall cell number/density is constant throughout these oscillations (Klevecz *et al.*, 2004; Tu *et al.*, 2005), which becomes important for our mathematical model.

### Defining the two states and apparent bistability

If these data are more grossly binned into groups, there appears to be ~2 effective equilibrium states in this system. If binned based on the gene/metabolic patterns, there is the oxidative phase (high oxygen consumption) closely coupled to growth, immediately followed by the reductive mitotic phase, which depends upon (and follows directly from), the oxidative phase. Indeed, experimental data suggest that these two steps, the growth and proliferation steps, are irreversibly coupled (Laxman *et al.*, 2010). This can therefore be conceived as one bin, representing a “growth” state. The extended, low oxygen consumption phase where there is a long, steady build-up of resources, can be viewed as a second bin. Both these states or bins appear to be somewhat stable, contained systems, with what appears to be a transition or inflection point leading to a committed switch to the other state. Thus, there appears to be an *apparent* cellular state bistability occurring during these oscillations in oxygen consumption. The stable, low oxygen consumption phase can therefore be practically envisioned as representing the non-dividing, “quiescent” state (Q), while the rapid increase in oxygen consumption followed by the reduction in oxygen consumption phase represents the “growth” state (G) (Figure 1E). Considering this, our objective was to build a mathematical model that conceptualized the oscillations in oxygen consumption as oscillations between these two (Q and G) states.

For this, we first needed to define what plausible, broad scenarios this YMC system might fit into. We therefore considered the currently accepted explanations for commonly observed cellular heterogeneity within clonal populations. Many microbial cells at high cell densities put out “quorum/alarmone” molecules that affect the entire population, and lead to collective behavior along with heterogeneity (Miller and Bassler, 2001; Schauder *et al.*, 2001; Whitehead *et al.*, 2001; Zhu *et al.*, 2003; Chen *et al.*, 2004; Farewell *et al.*, 2005; Srivatsan and Wang, 2008). Other possibilities emerge from metabolic resource sharing, seen widely in systems ranging from microbial populations to cancer cells (Veening *et al.*, 2008; Cairns *et al.*, 2011; Campbell *et al.*, 2015, 2016). This extends to regulation at the levels of metabolic specialization and stochastic gene expression resulting in phenotypic heterogeneity (Avery, 2006; Ibanez *et al.*, 2013; Holland *et al.*, 2014; Ackermann, 2015; Sumner and Avery, 2017). From within this range of possibilities, we envisaged three general scenarios that could result in the type of oscillations (Q <-> G) seen in the YMC, and could make biological sense (Figure 1F): there could be the production and secretion of a resource by a sub-population of cells (“feeders”), which is taken up by other cells that will go on to divide; (ii) there could be the secretion and accumulation of a metabolite that is sensed and taken up by only some cells (but is not consumed); (iii) there is a build up of a metabolite, which is consumed by the cells at some threshold concentration (Figure 1F). Starting from these scenarios, we built simple models to test which one could create an oscillatory system between the two states, which can come from an apparent bistability in the system.

**Figure 1:**
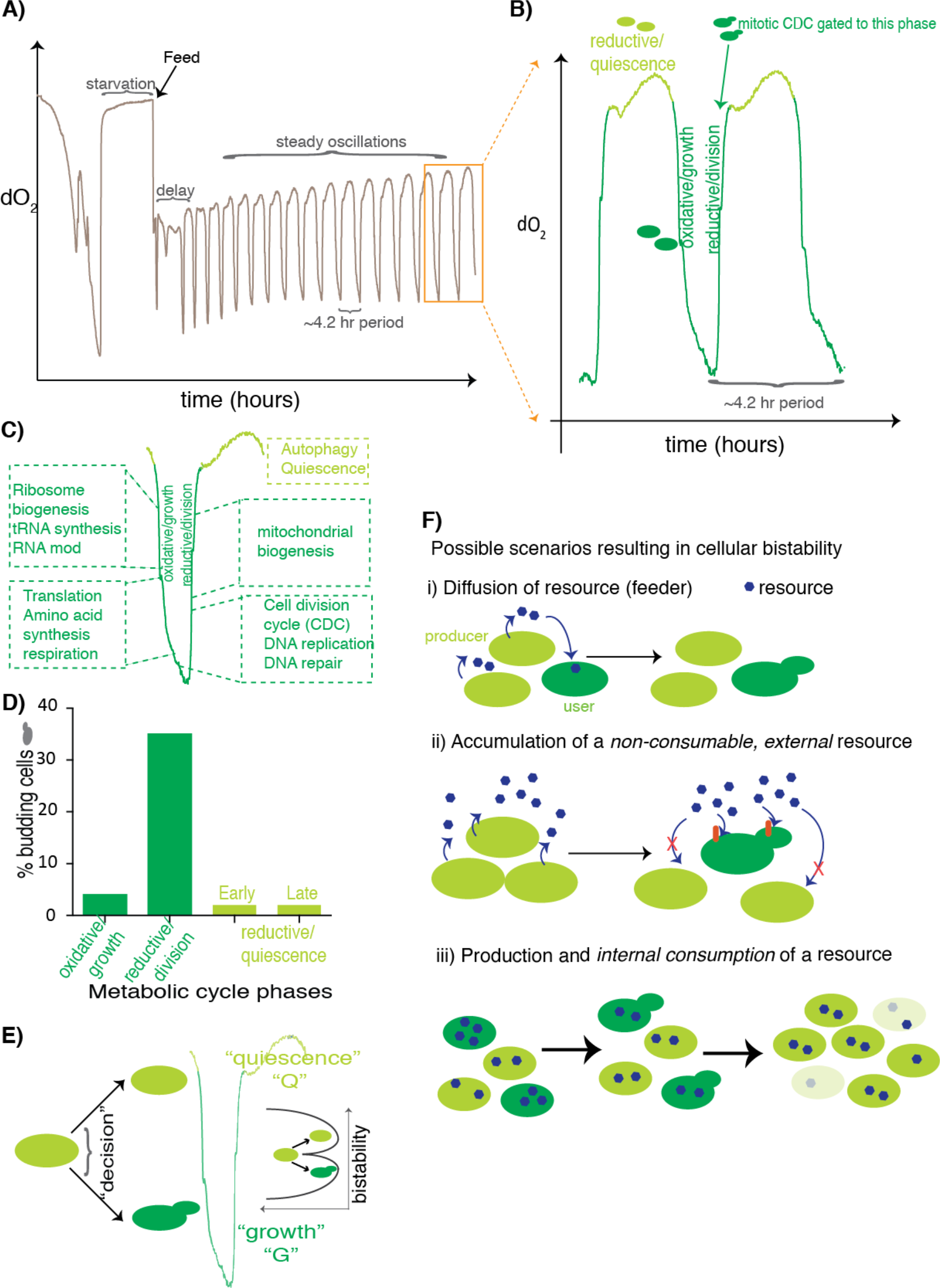
Apparent two-state bistability during Yeast Metabolic Cycles. A) A representative YMC, indicating stable oscillations in oxygen consumption (based on dissolved oxygen dO2) in yeast cultures, reflecting the yeast metabolic cycle. Note that the stable oscillations are driven by restricted feeding. B) A more detailed illustration of each oscillation cycle, also indicating the phases of the YMC. C) Functional outputs based on gene expression studies (from (33)), which clearly define the oxygen consumption phases of the YMC into a general “growth/proliferation” phase, and a “quiescence” phase. D) Observed cell division during the YMC. Cell division is tightly gated to a narrow window of the YMC. Note that only a fraction of cells, and not all cells, divide during this window of each cycle. E) Reducing the oxygen consumption (dO2) oscillation into a two-state (Q state and G state) system. The apparent bistability is also illustrated. F) Plausible biological scenarios that could result in an oscillation between Q and G states, based on observed phenomena. These scenarios are considered for building the model.

### A “push-pull” model, requiring communication between the Q state, G state and the resource, produces oscillatory behavior

#### Model framework for a two-state yeast population

In order to model such a two-state population of cells, the variables to consider would be the following: (a) The number of cells in the quiescent state and in the growth state, (b) some indicator of resource availability (dependent on the accumulation and consumption of the resource) which could modulate the switching rate between Q and G states, and the growth rate.

Thus, using this framework, we build the following equations that can describe the dynamics of a two-state population of yeast cells in a well-mixed system:

“Change in Q population over time”:

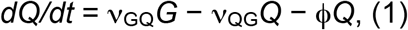

“Change in G population over time”: 
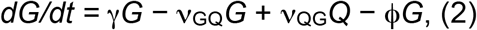
where Q*(t)* is the number of cells in the quiescent state at time *t*, *G(t)* the number of cells in the growing/dividing state, each ν represents a switching rate, ϕ*(t)* is the chemostat outflux rate (which could vary with time), and γ is the growth rate of cells in the growing/dividing state. If we further assume that the chemostat is working in a mode that maintains the total population (or density) of cells at some constant level, i.e., the outflux from the chemostat balances the growth of cells at all times, this means ϕ*(t)* = γ*G/(G + Q).* In this case, the population dynamics can be described by a single equation: 
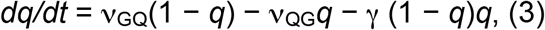
where *q* ≡ *Q/(G + Q)* is the fraction of cells in the quiescent state.

Next, we assume that the cells contain some ‘resource’ that they require for growth, without making any further assumptions about the resource. Let *a(t)* denote the concentration per cell of this resource at time *t*, and let σ denote the rate at which additional amounts of this resource enter each cell from the surroundings (where the resource is replenished due to the influx of fresh medium into the chemostat). *a* is depleted both by dilution due to the outflux (at a rate γ(1−*q*) as explained above), as well as by consumption by growing cells (this rate is also proportional to γ(1−*q*), which is the net rate of production of new cells). The dynamics of this resource can thus be described by the equation:

“Change in resource over time”:

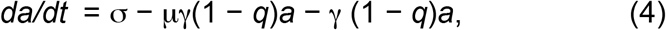
where μ is a proportionality constant that sets just how much resource is consumed by a growing cell, compared to the amount that is depleted by dilution.

In writing equations 3 and 4, we have assumed that all cells have the same amount of this internal resource *a*. A less restrictive assumption that still gives the same equation is to assume that *a* represents the average concentration of the resource across the population of cells, but that the distribution of resource levels is similar for Q and G cells. Further, the same equations also model the case where the resource is not an intracellular one, but an extracellular one - σ then is just reinterpreted as the rate at which the resource is added to the extracellular medium either by an external feed or by secretion of the resource by the cells themselves (e.g., by making σ dependent on *q*).

By choosing which of the parameters in the above equations are zero or non-zero, and how they depend on *q* and/or a, this framework can be used to model a variety of scenarios, which subsume the broad, biological scenarios illustrated in Figure 1E.

These mathematically distinct scenarios are described below (and illustrated in Figures 2A and 2B):

1. A sub-population of feeder cells (in the Q state) secrete a resource that is sensed by other cells that can grow and divide (G state); resource accumulation σ increases with *q*. Such a scenario can be modelled with the G cells either consuming the resource (μ ≠ 0), or only sensing but not consuming the resource (μ = 0) in the processes of growing/dividing. The growth rate in the G state may be a constant, or may depend on the level of the resource (e.g., γ proportional to *a*). There are three sub-scenarios for how cells may switch between the two states:

a. There is no switching between Q and G states (ν_QG_ and ν_GQ_ both zero).
b. There is random switching between Q and G states (ν_QG_ and/or ν_GQ_ are non-zero constants).
c. Switching between Q and G states is dependent on cell density and/or the resource level (ν_QG_ and ν_GQ_ both functions of *q* and/or *a*).
d. All cells produce and secrete a resource that is sensed only by a sub-population of (G) cells that can grow and divide, i.e., *σ* is a constant. As in scenario 1, the G cells may or may not consume the resource, the growth rate in the G state may or may not depend on the level of the resource, and there are three sub-scenarios for how cells may switch between the two states: no switching, random switching or density/resource dependent switching.
e. There is a build up of a resource, which is directly supplied from outside into the chemostat medium (σ is a constant). This metabolite is sensed or consumed by the G cells when they grow/divide. Again, the growth rate in the G state may or may not depend on the level of the resource and switching may work in one of three ways: none, random or density/resource dependent switching.

While scenarios (2) and (3) may appear mechanistically very different, they are in fact mathematically no different from each other; both result in a constant production of the resource (Figure 2B). Hence, we need not distinguish between these two. Testing all the scenarios above, using equations 3 and 4, we show in the next section that oscillations are not possible in the absence of switching, or even with random switching, when there is no substantial time delay between resource utilization and division events (as assumed in writing equations 3 and 4). Thus, scenarios 1c, 2c and 3c are the only possibilities left that give oscillations (Figure 2C). This means that the switching between Q and G states is a stochastic event, but with a probability that depends on the resource level, and/or the density of cells in the Q or G state, implying some form of communication between the resource, the cells in the Q state and the cells in the G state.

**Figure 2:**
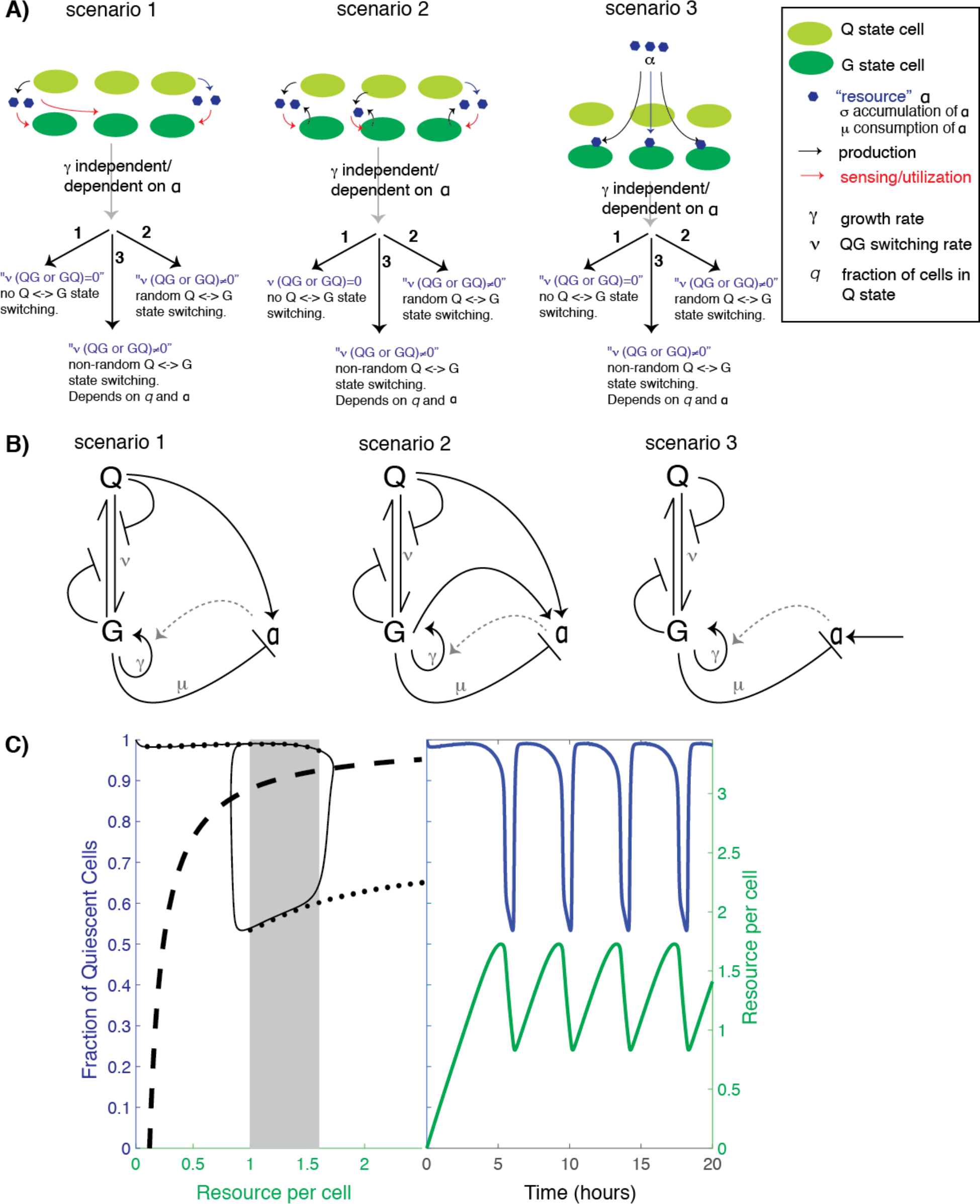
A “push-pull” model for oscillations arising from an underlying bistability between Q and G states. A) A range of biologically plausible scenarios from Figure 1F, now broken down into precise categories, where parameters affecting the rates of proliferation (g), switching between Q and G states (n), as well as consumption (m) and supply (s) of the resource are included. The variations in these parameters are used to build and test our model. B) Schematic illustration of Figure 1A, indicating feedback loops and parameters considered, to test for possible oscillations between Q and G states. For clarity, potential feedback loops caused by the parameters being dependent on the resource a are not shown, but are included in our models. C) A hysteretic oscillator, based on switching between Q and G states, a required communication between Q, G and the resource, and the oscillation of the amounts of resource itself that controls the Q<->G transitions (see the Methods for the parameter values that that produce this dynamics). In the left panel: The thin black curve shows the path traced by the oscillation in the q-a plane, the thick dashed line is the curve along which production of resource exactly balances consumption/dilution, and the solid black dots trace the high and low branches of the steady state q levels when the resource level is held constant (the grey rectangle indicates the region of bistability). In the right panel: blue and green curves show, respectively, the fraction of quiescent cells and the resource level as a function of time.

#### Some necessary conditions for oscillations

Within the framework of our model we can show that a density-dependent switching rate is necessary to get oscillations.

i. *No oscillations in the absence of switching:* When both ν_QG_ and ν_GQ_ are zero, then equation 3 above becomes:

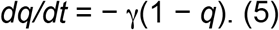 As long as γ is always positive, irrespective of its dependence on a, this has only one stable steady-state solution, *q* = 0, because the rate of change of *q* is always negative. And this is globally stable, i.e., every initial value of *q* (except *q* = 1) will flow to *q* = 0. The *q* = 1 state is an unstable steady state, i.e., any fluctuations away from it, however small, will result in the system moving to *q* = 0. Thus, there can be no oscillations in the absence of switching.
ii. *No oscillations with constant parameters:* When all the parameters in equations 3 and 4 are constants, independent of *q* and *a*, then no oscillations are possible because eq. 3 becomes independent of eq. 4, and therefore, being a one dimensional ordinary differential equation without explicit time-dependence, cannot show oscillations (an oscillation in *q* requires that *dq/dt* take both positive and negative values for the same value of q, for at least within some range of *q*, and this is not possible for a 1D ordinary differential equation).
iii. *No oscillations for random (density-independent) switching:* A less restrictive assumption is that ν_GQ_ and ν_QG_ are constants (which includes zero - we‘ve already examined the case where both are zero above), but γ and σ may be functions of *q* and/or *a*. In the scenarios we examine, γ may be an increasing function of *a* (all scenarios), while σ may be an increasing function of *q* (scenario 1). In this situation, the dependence of each variable on the other is ‘monotonic’ (*dq/dt* is a decreasing function of *a*, while *da/dt* is an increasing function of *q*). Equations with such monotonic dependencies have been studied mathematically in detail (Pigolotti *et al.*, 2007; Tiana *et al.*, 2007), which show explicitly that when such a coupled set of equations has only two variables (here, *q* and *a*), then sustained oscillations are not possible. Intuitively, there is not enough time delay in such a small two-leg feedback loop to destabilise the overall negative feedback that pulls the variables into a single stable steady-state value.

#### Hysteretic oscillator based on the two-state model

Apart from there being broadly two states, a second crucial observation from the experiments is that there is a distinct separation of timescales. The transitions from a situation where almost 100% of cells are in the Q state to one where 30-40% are in the G state, and vice versa, are very rapid. Whereas, between these two transitions the dynamics proceeds on much slower timescales. A simple way to obtain such a two-timescale oscillator from this two-state model uses the strategy of ‘frustrated bistability’ previously suggested by (Krishna *et al.*, 2009). It requires three ingredients: (1) a negative feedback loop between *q* and *a*, (2) bistability in *q* in the absence of the feedback, and (3) the assumption that changes in *q* happen on a relatively fast timescale compared to changes in *a*. While the first can be achieved in several ways, the two simplest, biologically plausible, scenarios are where growing cells consume a resource *a* and: (i) the growth rate γ is proportional to the resource *a*; or, (ii) one or both switching rates depend on *a* such that the net switching rate from G to Q decreases with a. However, the third requirement of separation of timescales means that the switching rates must be at least several-fold higher than γ and σ. This means that the term γ*q(1-q)* in equation 3 is practically negligible and hence the dependence of γ on *a*, or lack of it, would have little effect. We therefore concentrate on the case (ii) where the switching rates depend on *a* to implement the negative feedback, and for simplicity keep γ independent of *a*.

Bistability in *q* in the absence of the feedback implies that when *a* is kept fixed, for some range of *a* values, equation 3 should allow two stable steady state levels of q, one lower and one higher. This is shown in Figure 2C left panel, where one can see the high and low ‘branches’ traced by the solid black circles - every point on these branches is a stable steady state *q* can attain for the corresponding *a* value, using a version of equation 3 derived from scenario 3 in Figure 2B (see Methods for the full equation). When the resource *a* is sufficiently small, then there is only one high steady-state level possible for *q*. Similarly, when *a* is sufficiently large, there is only one low steady-state possible. However, for intermediate values of a, the system exhibits bistability and both low and high steady-state levels co-exist. In this bistable region, which steady-state level *q* attains depends on where it started (i.e., its ‘initial condition’). Importantly, in these oscillations, the system exhibits a ‘memory’ (or a ‘hysteresis’) - the steady-state level that *q* eventually settles into depends on the history of the system.

When there exists such bistability, then one can get oscillations from the system described by equations 3 and 4, provided the switching rates are a few-fold faster than the rates of consumption and accumulation of the resource (Krishna *et al.*, 2009), as follows: when *q* is high, *a* increases due to lack of consumption, so the system creeps along the high branch in Figure 2C left panel (see the trajectory traced by the thin black line) until it hits the edge of the bistable region. At that point, cells start switching to the G state, which happens relatively rapidly due to the separation of timescales. Thus, the trajectory “falls off” the edge down to the low branch. On the low branch, with more G cells, the now increased consumption of the resource causes *a* to start decreasing, leading to the system creeping down along the low branch. When the system reaches the left edge of the bistability, the trajectory jumps up to the high branch as cells rapidly switch to the Q state. For a range of parameter values, this settles into a stable oscillation, as shown in Figure 2C right panel, which shows how *q* and *a* vary with time as one follows the black trajectory in Figure 2C left panel.

For this kind of oscillation, as we have demonstrated in the previous section, ν_QG_ and/or ν_GQ_ must necessarily be functions of q, not constants independent of q. This can be interpreted as a form of ‘*quorum/cell number sensing*’ - implying some form of cell-cell communication (or a cell density dependent phenomenon). More specifically, we find that choosing either v_QG_ to be a decreasing step-function of *q* (as in Figure 2C), or ν_GQ_ to be an increasing step-function of *q* (see Supplemental Figure S1) is sufficient to produce frustrated bistability. Other shapes that we have not explored may also produce bistability, and hence oscillations. However, our purpose here is not to find the ‘best-fit’ model, but rather to demonstrate the basic ingredients which are sufficient to produce hysteretic oscillations that are similar to the experimental observations. The requirement for ν_QG_ to be a decreasing step-function of *q*, or Dν_GQ_ to be an increasing step-function of *q*, is basically a requirement for a “push-pull” mechanism - the more the Q cells, the more other Q cells get pulled to remain in that state, and the more G cells get pushed to switch away from their state, and vice versa. Irrespective of the precise molecular means by which this is achieved, cell-cell communication is a necessary ingredient for implementing such a push-pull mechanism.

#### Possible variations in the shape of the oscillations

From our gross model explained in Figure 2, we obtain predictable oscillations with a specific pattern. The model oscillations exhibit a fast drop in *q* when exiting the predominantly quiescent phase, followed by a slow(er) drop, and then a rapid rise back to a high *q* level. Experimentally however, a few variations within the general oscillation patterns are known to occur, depending upon the strain background (Burnetti *et al.*, 2015). In the CEN.PK strain (our major reference system, from where the gene and metabolite oscillation datasets were obtained (Tu *et al.*, 2005, 2007; Mohler *et al.*, 2008)) dO2 levels (which we equate with q) show a fast drop, a slow further drop, and then a rapid rise (Figure 3A scenario (i)). However, as comprehensively described in (Burnetti *et al.*, 2015), three other variations have been extensively documented. Following a fast drop in dO2 levels, some strains then show a slower drop followed by a more extended low dO2 phase (bump), and a fast rise in dO2 (Figure 3A, scenario (ii)). Other strains show an overall fast drop in dO2, an extended low dO2 phase and bump, and a fast rise (Figure 3A, scenario (iii)), or a fast drop in dO2 (increased oxygen consumption), followed by a slower, extended rise in dO2 (Figure 3A, scenario (iv)).

#### Can our model explain this small diversity of shapes seen during the overall drop and rise in oxygen concentrations?

In the model, the shape observed depends on the shapes of the two branches of *q* steady-states (solid black circles in Figure 2C, left panel). Because the lower branch starts at a *q* value of around 0.5 and then increases as *a* increases, therefore there is a slow drop in *q* after the fast drop. To produce the experimental dO2 oscillations in other yeast strains (as shown in Figure 3A), the lower branch must have a different shape.

For example, for strains which show a slow increase after the first rapid decrease of *q*, the low *q* branch must *decrease* as *a* increases. Similarly, the other waveforms would involve other shapes of the lower or higher branches. In Figure 3C we show that simple changes in the dependence of the switching rate ν_QG_ on *a* produce different waveforms for the oscillations. Here we’ve shown how to get different shapes of the low-*q* phase of the oscillation by manipulating the lower branch of the bistability - changes to the high-*q* phase could similarly be easily made by manipulating the upper branch. The main point is that the shape of the waveform is primarily determined by the shape of the bistability branches, which in turn are determined by how ν_QG_ and ν_GQ_ depend on *q* and *a*. Thus, our model predicts that these switching rates are what must vary between strains that show different oscillation waveforms.

**Figure 3:**
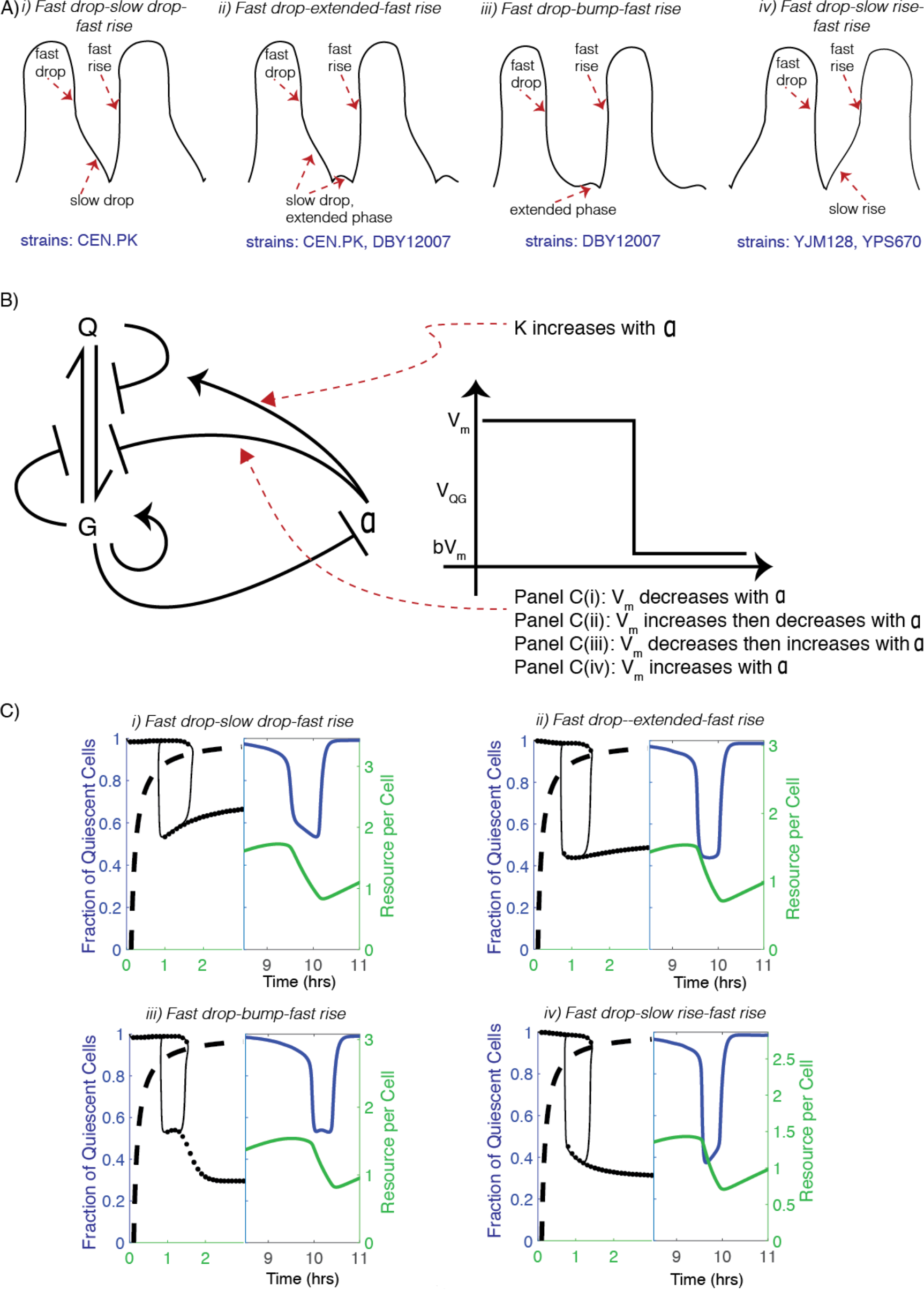
Diverse waveforms in the oscillations: experimentally observed and model predictions. A) Experimentally observed patterns of oscillations in dissolved oxygen/ oxygen consumption, which is dependent on yeast strain backgrounds and chemostat growth conditions. B) Altering the communication loops between Q, G and a, to change the overall oscillation waveform. Here g (growth rate) is constant and n_QG_ is a decreasing step-function of q. To obtain different waveforms, we vary the way the step function parameters n_m_ depends on a. C) Predicted oscillation patterns from the model (as altered described in panel B). The illustrated panels cover the range of waveforms observed experimentally in panel A. (i) same as Fig 2C; n_m_ decreases with a. (ii) n_m_ first increases then decreases with a. (iii) n_m_ first decreases and then increases with a. (iv) n_m_ increases with a. Additionally, in all four cases, K increases with a and other parameters have been chosen so that the time period of oscillations is close to 4 hours (see Methods for the full equations, with parameter values, for each case).

#### Predicting oscillatory outcomes based on resource availability

We have used scenario 3c (from Figure 2) to produce oscillations in Figures 2 and 3 above. We reiterate that mathematically scenarios 2 and 3 are the same, so scenario 2c can produce exactly the same oscillations. Further, we also find that scenario 1c (where the resource is not supplied externally, but produced/secreted by only the Q cells) is also capable of producing similar oscillations, based on highly constrained choices for how the production rate of the resource (σ) depends on *q* and *a* (see Supplemental Figure S2). Thus, while scenarios 3c and 2c are identical, all three scenarios, 1c, 2c, 3c, with appropriate choices for how the switching rates, and production and consumption depend on the resource and fraction of quiescent cells, are sufficient to explain the YMC oscillations. Scenario 2c and 3c are largely indistinguishable, and both appear biologically most plausible. Given our experimental understanding of the YMC (and the need for a consumable resource, glucose, to control the oscillations), we think scenario 3c is most likely (and we will explore this further in a subsequent section).

### Breakdown of the oscillations

In Figures 2 and 3 above, we have chosen the particular “default” values of each of the model parameters such that the oscillation period became approximately 4 hours, to match the experimental observations in Figure 1. Of course, varying these parameter values changes the time period, and for large enough variation the oscillation may also disappear. Our model predicts how the oscillation shape and period will vary, and when oscillations will break down, in response to experimentally tunable parameters. For instance, Figure 4A shows how the oscillations change as the resource production rate, σ, is varied around its default value, for the same equations that produced the oscillations in Figures 2 and 3. When *σ* is decreased below the default value, the oscillation period initially increases, with more time being spent in the high-q phase. For low enough σ, the model exhibits damped oscillations, and then as *σ* is lowered further, the model exhibits the absence of oscillations, with *q* settling into a high steady-state value (see Figure 4A, and also Supplemental Figure S3 for more such plots). When *σ* is increased from its default value, we again find that the period initially decreases, with less time being spent in the high-*q* phase. We are able to produce oscillations having a time period as low as ~2.5 hours (see Figure 4A(iii)). When *σ* is increased beyond this, the oscillation period starts increasing again, and the low-q phase of the oscillation starts becoming pronounced (see Supplemental Figure S3). Eventually, the oscillations disappear, with *q* settling into a (relatively) low steady-state value. These predictions largely mirror known experimental observations, where decreasing or increasing feed rate (at these scales) control oscillations similarly.

The resource production rate σ is a parameter that can be tuned relatively easily in a chemostat by controlling the amount of fresh glucose or ethanol being supplied per unit time. However, another parameter that may be tunable by genetic modifications is γ, the growth rate of cells when they are in the G state. Figure 4B shows how the oscillations vary as γ is varied. The results are qualitatively similar but inverse to what was observed with σ variation - an increase in γ from the default value results in an increasing period, damped oscillations and eventually no oscillations, while a decrease first results in a decrease of period, then a distorted shape and increasing period (see Supplemental Figure S4 for more such plots).

The location of the dashed black lines in Figure 4 (left panels) help to understand this behaviour. Each dashed line traces the *q* and *a* values where resource production exactly balances resource consumption/dilution. To the right of the line the production is less than the consumption so the resource must decrease, and vice versa to the left of the line. The closer one is to the dashed line, the slower the rate of change of a. As explained in (Krishna *et al.*, 2009), oscillations occur only when this dashed black line passes between the upper and lower bistable branches (solid black circles) - because then the resource keeps increasing (decreasing) when it reaches the end of the high (low) branch making the trajectory “fall off the edge” and continue the oscillation. When σ is decreased the dashed black line shifts leftward in the plot, coming closer to the high *q* branch which causes the oscillating trajectory to spend more and more time on the high *q* branch (because it is closer to the dashed line and so the resource accumulates slower). Eventually, as the dashed line just touches the high *q* branch, the time period of oscillations increases to infinity (logarithmically - see Supplemental Figure S5). For σ values lower than this critical value there is no sustained oscillation and the system settles into a steady-state on the high branch at the point where it crosses the dashed line. A similar behaviour happens as σ is increased and the dashed line comes closer to the lower branch, with the only difference being that the oscillating trajectory spends more time at lower *q* values.

A universal feature of the YMC oscillations seen in diverse yeast strains is that the time period of the oscillations decreases with an increase in the dilution/supply rate in the chemostat. The time period appears to be dominated by the time spent in the high-q phase, which also increases with dilution/supply rate, whereas the time spent in the low-*q* phase is less and *decreases* slightly with increase in the dilution/supply rate (described in (Burnetti *et al.*, 2015)). As described above, in our model, we find that as we vary σ or γ, there are two regimes. In one the time period is dominated by the high-q phase, and the behaviour matches the above experimental observations (see Supplemental Figure S5). However, there is also another regime, where the time period is dominated by the low-q phase. Our model therefore predicts that: (i) the observed YMC oscillations are closer to the lower end of the *σ* range that produces oscillations, so one should be able to increase *σ* more than decrease it before breaking the oscillations, and (ii) if one increased *σ* enough while remaining in the oscillatory regime one should observe low-*q* dominated oscillations such as those shown in Supplemental Figure S3. These are both testable predictions of our model.

**Figure 4:**
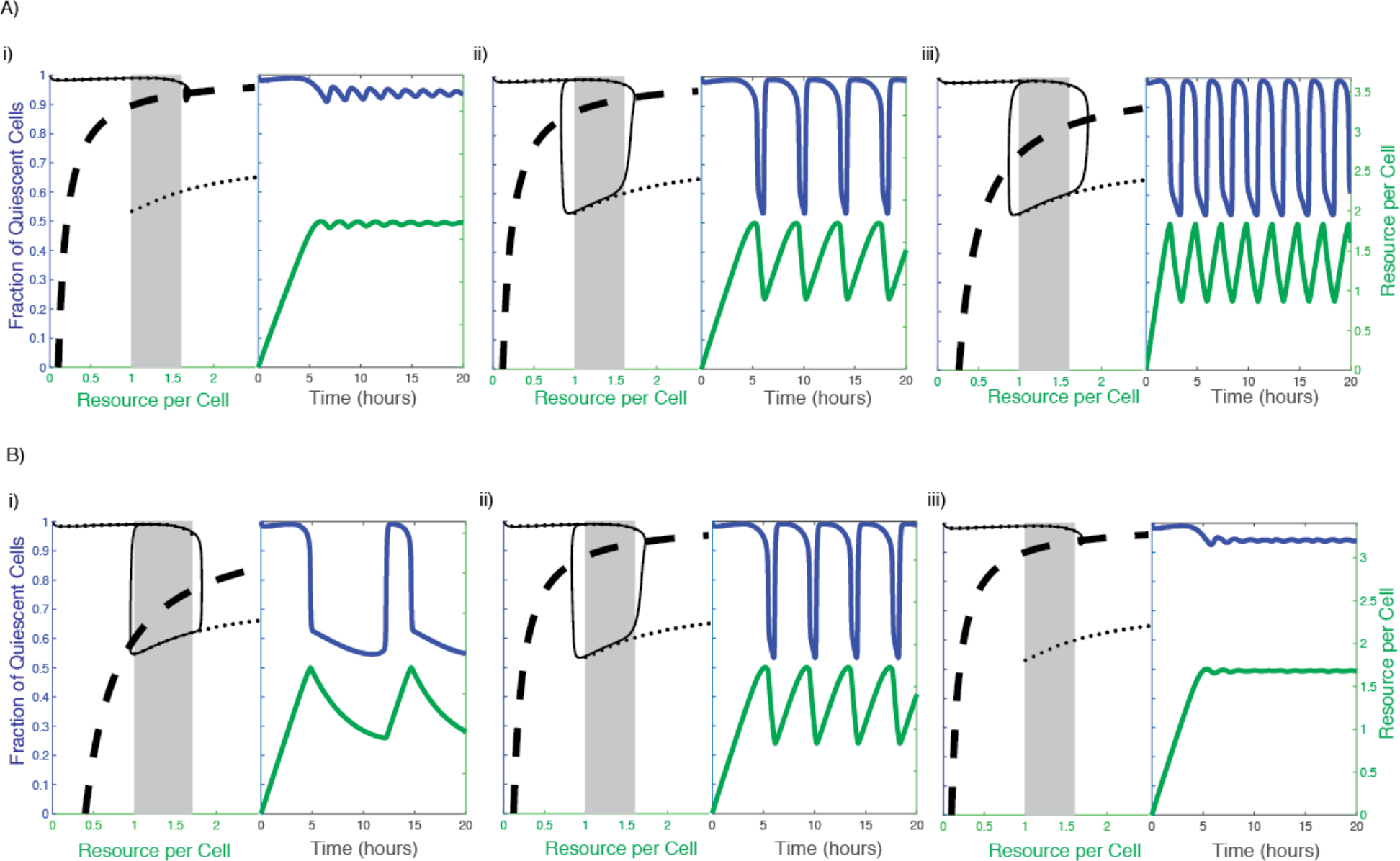
Breakdown of oscillations. A) Varying the rate of production of resource σ. (i) σ = 0.346 hr, (ii) σ = 0.400 hr (default parameters, same as Figure 3), (iii) σ = 0.866 hr. B) Varying the growth rate of cells γ. (i) γ = 0.500 hr, (ii) γ = 1.665 hr (default parameters, same as Figure 3), (iii) γ = 2.000 hr. Equations used, and other parameter values, are the same as those that produced Figs 2C and 3C(i).

### Acetyl-CoA and NADPH satisfy the requirements of the consumable resource that controls oscillations between Q and G states

Based on our model, the metabolic resource oscillates with a unique pattern, and this drives the oscillation between the Q and G states. From the model, some resource builds up within the cell, and is highest at the point of commitment to the switch to the G state (Figure 5A). It is then rapidly consumed/eliminated to fall below a certain threshold, resetting the oscillation, after which the cycle of building up for consumption resumes. When superimposed to the actual YMC phases (and the Q to G switch), this build-up of the resource would necessitate its highest amounts at the beginning of the phase where cells commit to entering high oxygen consumption (Figure 5A). We note that these features of the resource oscillation are a very robust prediction of our model. Across all the oscillations in Figs 2-5 we see the same behaviour, and we would see this for any parameter choice that gives oscillations because this behaviour depends only on our assumption that the resource is consumed by growing/dividing cells and not by quiescent cells. Therefore, according to our model, in order for any metabolite to be the resource that controls the oscillation between the two states, this molecule must fully satisfy the above criteria. Furthermore, for completing this switch to the G state, the metabolite must be able to drive all the downstream biological events for growth. So do any central metabolites satisfy these requirements, and could therefore be the internal resource that controls these Q-G oscillations?

**Figure 5:**
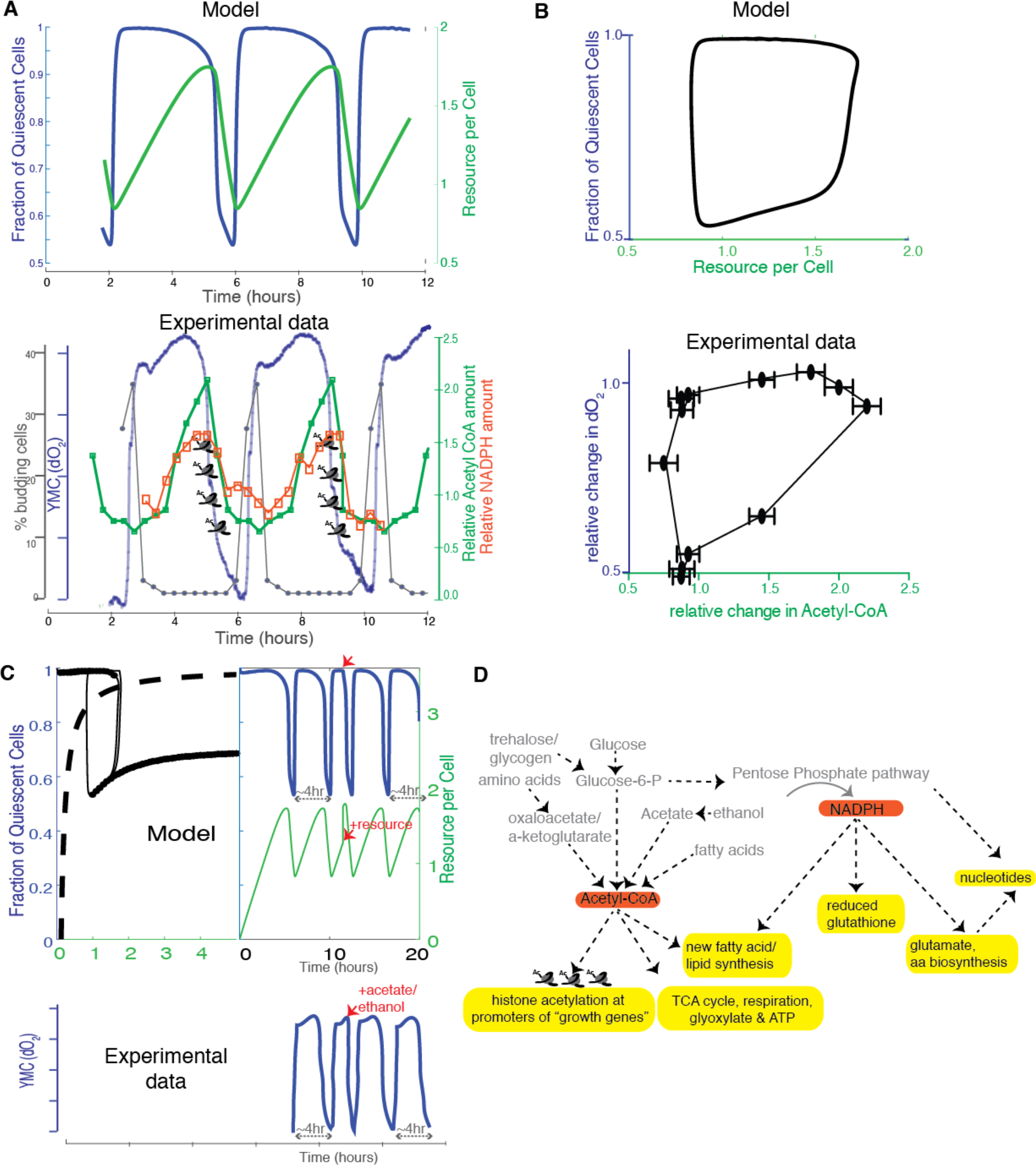
Acetyl-CoA satisfies the requirements for the metabolic resource controlling the Q and G oscillations. A) Predicted pattern of oscillation of the resource, during the Q and G oscillations, based on the model (top panel, same oscillations as Figure 2C), and experimentally observed oscillations of acetyl-CoA and NADPH during the dO2 oscillations (bottom panel). B) Predicted phase portrait of the the fraction of quiescent cells vs the resource per cell based on the model (top panel), and experimentally observed oscillations in dO2 and acetyl-CoA. C) Predicted effect on the oscillation waveforms and the Q and G states, when a bolus of the resource is added to cells in the Q state (see Methods for details), vs experimentally observed data on oxygen consumption when a resource, acetate (the trace is similar with for resources like ethanol, acetate, acetaldehyde, glucose) is added to cells in the low oxygen consumption phase. Supplemental Fig S6 shows how the response varies as the time of adding the bolus is varied. D) Acetyl-CoA as a central regulator of a switch to the growth (G) state. The schematic illustrates a cascade of biological processes leading to growth that acetyl-CoA amounts regulate (coupled with coincident, required NADPH utilization). Note that all resources that reset the oscillations, as indicated in (C) are utilized after they are converted to acetyl-CoA.

Comprehensive datasets of 50-100 oscillating metabolites in the YMC exist (Murray *et al.*, 2007; Tu *et al.*, 2007; Mohler *et al.*, 2008). From these studies, the oscillations of only two metabolites, acetyl-CoA and NADPH, fully fit the criteria demanded by our model. The acetyl-CoA and NADPH oscillations as a function of the metabolic cycle, and transitions between the Q and G state are shown in Figure 5A and 5B. The oscillations of acetyl-CoA during the YMC almost perfectly superimposes with the oscillation pattern of the hypothetical metabolic resource predicted by the model (Figure 5A). We plotted phase diagrams of the fraction of quiescent cells vs the amount of resource in the cell (from the model), and also plotted phase diagrams from experimental data for the dO2 oscillations plotted against acetyl-CoA amounts (Figure 5B). The two phase diagrams (from the model, and from experiments) strikingly resemble each other (Figure 5B). This is despite the fact that the experimental data for acetyl-CoA is of low resolution, with only a few sampling/time points covered, and also only reflects overall (bulk population) measurements of acetyl-CoA, suggesting that the actual phase diagram might be even more similar. Thus the model appears to capture key universal features of these yeast oscillations, including the point of exit from low oxygen consumption (Q) to high oxygen consumption and back (G), and the parameters important in the waveform (i.e. the low and high oxygen consumption phases are important, while the precise form of the dip and increase in dissolved oxygen may not be so). The model also supports an inference that the acetyl-CoA oscillations are sufficient to explain the bistability between the Q and G states, and retains the hysteresis component.

Using our model, we next simulated what would happen if the resource was increased to just above the threshold level, at a different time. In our model, during normal oscillations, the amount of resource steadily increases, while the cells are in the Q state. In our simulation, we provided a single bolus of the resource, while cells were in the Q state (Figure 5C). We observed a predictable, sharp exit from the Q state and entry into the G state (Figure 5C), effectively resetting the oscillation, which then continued and restored itself to the normal, ~4 hour period in the next cycle. We compared this to available experimental data, where oscillations have been reset by adding a bolus of an external agent, typically glucose, acetate, acetaldehyde or ethanol (Murray *et al.*, 2003; Klevecz *et al.*, 2004; Tu *et al.*, 2005; Cai *et al.*, 2011). All these agents show near identical patterns of resetting of oscillations (exit from Q and entry into G), and a representative figure (for acetate addition) is shown in Figure 5C. Here, cells exit the low oxygen consumption phase and enter and exit the high oxygen consumption phase, and subsequently quickly restore normal (in this case ~4 hr) oscillations. This simulation can be done in any part of the oscillation, and whenever most cells are in Q, adding a bolus of the resource similarly resets the oscillation (Supplemental Figure S6). Also notably, adding this resource when cells have switched to the G state does not alter the oscillations much (Supplemental Figure S6), which is also something widely established in experiments. Thus, the oscillations predicted by the model very closely recapitulates the patterns of oscillations observed in experiments, how the central, controlling resource might oscillate, and how the oscillations are affected upon perturbing the resource. This strongly suggests that the threshold amounts of the resource are sufficient to set the oscillations and switching between Q and G states.

Multiple lines of experimental data suggest that these two metabolites, acetyl-CoA and NADPH, are key in controlling exit from quiescence and entry into growth (Tu *et al.*, 2007; Cai *et al.*, 2011; Cai and Tu, 2012; Machné and Murray, 2012; Shi and Tu, 2013, 2014; Mellor, 2016). Based on our knowledge of the metabolic prerequisites for entering growth, and known functional endpoints or outcomes of these two molecules (Figure 5D), we can now make a strong, parsimony based argument suggesting that oscillations in these two metabolites are sufficient to control oscillations between the Q and G state. Particularly, several lines of study suggest that the entry into growth (from quiescence) depends on carbon source utilization (Shi *et al.*, 2010; Cai *et al.*, 2011; Daignan-Fornier and Sagot, 2011; Laporte *et al.*, 2011). As pointed out earlier, studies from the yeast metabolic cycle show that the oscillations depend upon carbon sources (primarily glucose) (Klevecz *et al.*, 2004; Tu *et al.*, 2005), and oscillations can be reset (to enter the growth program) by adding acetate, acetaldehyde, etc. (Murray *et al.*, 2003; Tu *et al.*, 2005; Cai *et al.*, 2011). Notably, these carbon sources end up being converted directly to acetyl-CoA, and can only then be utilized (Figure 5D). Additionally, a growth program will require not just sufficient energy (ATP) to sustain the anabolic processes within it, but also activate a program boosting anabolic processes that lead to cell division, including making enough lipid moieties required for cell membranes and other constituents of a new cell. Notably, acetyl-CoA satisfies all these requirements in the following manner (Figure 5D): it directly enters the TCA cycle to generate ATP (Nelson, DL; Cox, 2017); it can be utilized for the biosynthesis of numerous cellular metabolites, including fatty acids, sterols, and amino acids (Nelson, DL; Cox, 2017); and directly regulates cell growth and ribosome biogenesis by the acetylation of histones at “growth promoting genes”, especially histones at ribosome subunit, tRNA and ribi genes, and activates their transcription by the SAGA complex (Cai *et al.*, 2011). The genes that breakdown storage carbohydrates (such as glycogen and trehalose) that produce acetyl-CoA all peak before the maximal acetyl-CoA concentration (Tu *et al.*, 2005; Kudlicki *et al.*, 2007). Finally, the exit from quiescence requires the liquidation of these storage carbohydrates (Shi *et al.*, 2010; Laporte *et al.*, 2011; Shi and Tu, 2013), and conversion to acetyl-CoA (and the subsequent gene expression program) (Shi and Tu, 2013). Perturbations in the ability to sense and utilize acetyl-CoA (particularly for the gene expression program) completely abolish oscillations (Cai *et al.*, 2011). Physiologically, this anabolic commitment also absolutely requires the process of reduction for anabolic biosynthesis, and this reductive capacity is supplied by NADPH (Nelson, DL; Cox, 2017) (Figure 5D). NADPH is primarily synthesized from the pentose phosphate pathway, which branches from this same central carbon network, and this NADPH will fuel the required reductive biosynthesis to make molecules required for anabolism (Figure 5D). Finally, genes encoding proteins that increase the synthesis of NADPH are similarly coincident with those that lead to the generation of acetyl-CoA, and disrupting NADPH production slightly results in a collapse of oscillations (Tu *et al.*, 2005, 2007). Relatedly, studies from the YMC show multiple other metabolite oscillations coupled to or dependent upon NADPH, although any hierarchical organization was not immediately apparent (Murray *et al.*, 2007). Without a necessary coupling of the two molecules, the overall process of entry to growth cannot be completed. There is substantial data, particularly from the studies of various cancers, to show the close coupling of acetyl-CoA and NADPH for growth (Heiden *et al.*, 2009), as well as direct evidence of acetyl-CoA promoting NADPH synthesis (Patra and Hay, 2014; Shan *et al.*, 2014). Summarizing, based on the pattern of oscillation of the resource predicted by our model, acetyl-CoA and NADPH (based on production and utilization) satisfy sufficiency requirements to be the molecules that control the Q-G state oscillations. Our model thus strongly supports an argument for oscillations in acetyl-CoA being sufficient to control Q-G state oscillations.

## Discussion

In this study, we present a simple frustrated bistability model to explain how the amounts of an internal metabolic resource can determine oscillations between a quiescent and growth state. For this, we relied on extensive data coming from the YMC, and represented the oscillations in dissolved oxygen (seen during YMCs) as a reflection of growth and quiescent states (Figure 1). Our model incorporates factors dependent on growth rate and amounts of the resource, as well as switching rates (between the G and Q states). Importantly, the model emphasizes a necessary communication between the cells in the quiescent state and the growth state, both of which interact with the metabolic resource during such transitions (Figure 2). Quiescent cells “push” cells in the growth state into quiescence, and “pull” other quiescent cells to remain quiescent, with the feedback requirements imposed by the resource being distinct and opposite for the Q and G states. Given this communication requirement between the Q and G states, our model suggests that such oscillations will eventually breakdown when the cell numbers are small and cells are no longer in contact with each other (something that has been experimentally observed (Laxman *et al.*, 2010)). This model also provides insight into understanding the “growth/division” rate of cells once committed to growth. While healthy debates continue on the rate of growth in a cell and stages of the cell cycle (Johnston *et al.*, 1977; Conlon and Raff, 2003; Jorgensen *et al.*, 2004; Brauer *et al.*, 2008; Goranov *et al.*, 2013). our model shows that it is sufficient for oscillations to have a fixed “growth rate” once the metabolic resource has crossed its threshold concentration, and triggered a committed growth program, after which the growth and division process is no longer dependent on available nutrients. This is also analogous to studies of the CDC, which are built around committed, “no return” steps that proceed at constant, predictable rates once committed to. In our model, because there is a timescale separation between growth and switching rates, making the growth rate dependent on the resource would make some quantitative difference to the rate of accumulation/consumption of the resource, but would leave the Q-G oscillations largely unchanged. Finally, using a parsimony based argument, we suggest that acetyl-CoA (along with NADPH) satisfies all requirements for the resource that drive these oscillations between the Q and G states (Figure 5). With acetyl-CoA as a resource, our model, which builds oscillations on an underlying hysteresis, reproduces universal features observed in these yeast metabolic oscillations, and provides a fairly simple sufficiency argument for how cells transition between Q and G states. We reiterate that our model only provides a paradigm to explain how the oscillations in an internal metabolic resource is sufficient to control oscillations between quiescent and growth
states. This allows for (but doesn’t include) other necessary elements in cells (e.g., unique gene transcription programs, or subsequent metabolic events that typically must follow), that may also be required to build a more detailed model for Q-G oscillations.

The kind of oscillator we have built falls under the class of “relaxation oscillators”, which have been used to model a very wide variety of phenomena, ranging from electronic oscillations to oscillating chemical reactions (Balthasar, 1926; Strogatz, 1994). These are a subset of several possible types of oscillators that arise in biological systems, and are especially relevant for the CDC (Novák and Tyson, 2008; Tsai *et al.*, 2008; Ferrell *et al.*, 2009; Ferrell, 2011). Relaxation oscillators typically involve the cyclic slow build-up of some quantity, like charge in a capacitor, until it reaches a threshold level which then triggers a “discharge” event, resulting in a rapid drop of the quantity. Thus, relaxation oscillators are often characterised by processes happening on two very different timescales, with the time period mainly determined by the slow process (Tyson *et al.*, 2003; Novák and Tyson, 2008; Tsai *et al.*, 2008; Ferrell *et al.*, 2009; Ferrell, 2011). This is why, in contrast to linear, harmonic oscillators, they can produce non-smooth oscillations like a square or sawtooth waveform. We note that the YMC oscillations show a clear signature of multiple timescales - in Figure 1 it is evident that the exit from quiescence (fast drop in dO2), as well as the re-entry into quiescence (fast rise in dO2), happen at much faster timescales than the other phases of the oscillation. In our relaxation oscillator model of the YMC, these differing timescales arise from the fact that the switching rates are an order of magnitude larger than the rates of production and consumption of the resource, and even the growth rate of the cells. The latter processes are therefore what determine the time period of the YMC. Within the class of relaxation oscillators, our models fall into a sub-class that depends on an underlying bistability, which is ‘frustrated’ (Krishna *et al.*, 2009). The bistability, and the resultant hysteresis, are what determine the threshold points at which the behaviour of the system rapidly switches between accumulating or consuming the metabolic resource. Interestingly, our model necessitates this strong hysteresis element within the Q and G state cells. The phenomenon of hysteresis has been extremely well studied (and established) particularly during many phases of the classical CDC, or proliferation cycle ((Pomerantz and McCloskey, 1990; Tyson and Novak, 2001; Solomon, 2003; Wei *et al.*, 2003; Angeli *et al.*, 2004; Han *et al.*, 2005; Ferrell *et al.*, 2009; Ferrell, 2011; Yao *et al.*, 2011) and many more). In contrast, a hysteresis phenomenon has not been extensively explored when cells transition between a growth state and an effective quiescence state. Yet, in such conditions where the transition between the two states is substantially determined by a metabolic oscillator, as seen in the YMC and several other studies from simple models like yeast, the hysteresis phenomenon is clearly revealed by our model. Given this, experimental studies can be designed to dissect the nature of this hysteresis phenomenon.

### General features emerging from the model to understand oscillations between quiescence and growth

Although our model is relatively simple, uses data from a fairly elementary system, and makes minimal assumptions, it does surprisingly well to constrain the possibilities for how transitions between quiescence and growth are regulated. The model successfully captures universally observed waveforms of oscillations, can reset the oscillations, can predict how the oscillations of a resource can control the two states, and can predict breakdown of oscillation fairly well, as observed in experiments. From the very large set of metabolites known to oscillate during the YMC (Tu *et al.*, 2007; Mohler *et al.*, 2008), our model constrains possibilities to a few, that oscillate in a way that can permit such a bistability to exist. From this, and consistent with extensive experimental data (discussed earlier, and in Figure 5), it is possible to make parsimonious arguments for acetyl-CoA (coincident with NADPH) as the metabolic resources controlling transitions from quiescence to growth, and *vice versa.* Our model helps differentiate this small set of metabolites from other metabolites that are important to maintain oscillations, but not initiate them (i.e. they may only allow the cell to continue in one state, or the other). For example, sulfur metabolism is critical to maintain oscillations (Murray *et al.*, 2003, 2007; Tu *et al.*, 2007). It is also essential for the completion of a growth program, post entry into the high oxygen consuming phase. But this metabolite peaks *after* acetyl-CoA in the YMC (Tu *et al.*, 2007), and can be viewed as a consequence of initiating a growth program, and also critical to sustain/complete this growth program, but not to initiate the oscillation. substantiating this explanation is the fact that sulfur metabolism is highly dependent upon the utilization of NADPH for reduction, and NADPH (as described earlier) is coincident with acetyl-CoA. A similar argument can be made for the sustained, high respiration seen in the YMC, which produces ATP that will be required to maintain the growth program once committed to by the cell. Separately, other studies have shown that “quiescent” cells can show metabolic oscillations without entry into the CDC (Slavov *et al.*, 2011). Here, these cells appear to show a commitment to the CDC during these oscillations, based on gene expression patterns (Slavov *et al.*, 2011). This can also be viewed through our interpretation of the commitment of cells to the CDC due to a central resource. Cells will commit to the CDC, which however may not be completed if a subsequent metabolic resource, normally dependent upon the central/controlling resource (predicted to be acetyl-CoA/NADPH here), becomes limiting. In other words, for a cell, usually if this committing resource is at the correct threshold, other resources should not be limiting unless artificially constrained in an experimental set-up. In (Slavov *et al.*, 2011), the limiting resource was phosphate, which typically should be available and not limiting, and be assimilated into nucleotides in an NADPH and acetyl-CoA dependent manner. If in a specific instance this resource becomes limiting, the cells would commit to the growth/CDC state, but will not be able to complete this, and will fall back into the Q state.

Our model provides a foundation to build new models to resolve other aspects observed during the YMCs. First, in each cycle of the YMC, a fraction of the cells exit quiescence and divide. It is not fully clear if the same cell divides in each cycle, or if a cell that has entered division in one cycle does not in the next, and so on. The decision to divide has been viewed as a stochastic, but irreversible step (Laxman *et al.*, 2010; Burnetti *et al.*, 2015). While our model as it stands cannot directly address these questions, the dependence of the oscillations on the build-up and utilization of a specific resource, allows the following argument to be made. First, the decision to divide in a cell would be purely made by the amount of resource (acetyl-CoA) that has been built up in the cell. Once acetyl-CoA reaches a certain threshold, the decision to divide is irreversible.

However, the build up of acetyl-CoA within an individual cell itself would be dependent on small differences in overall metabolic homeostasis (compared to its neighbor), and thus which cell reaches the threshold level first could be purely stochastic. Second, we may speculate that if a cell has reached this threshold level and then used up its resource during division, it is unlikely to be in a position to divide in the next round/next cycle, whereas a cell that had not reached the threshold level in the previous cycle would be best poised to divide instead. Our model does not take this into account, but it provides a framework within which one could model the entire distribution of cells in different Q/G states and with different levels of the resource. Despite the overall stochastic aspect of Q-G transitions, such models would be able to make testable predictions about the switching process even at the level of single cells. It is also apparent that this level of synchrony requires high cell density in the system.

Separately, most studies have noted that upon initiating feeding in the chemostat, there is a short period of tiny, non-robust oscillations. Based on our model, we would argue that this is a situation where the quiescent cells are all now building up just sufficient reserves of acetyl-CoA, within this stochastic process, and are starting to divide, but the unusual steady-state condition in the chemostat will eventually lead to stable oscillations.

Finally: Given the existing frameworks to describe Q-G state oscillations, our model is necessarily coarse grained, and is intended only to build a more rigorous conceptual framework within which to investigate the process of cells switching between quiescence and growth states. For instance, it is straightforward to extend our models, by adding space and diffusion processes, to account for scenarios where nutrients are not well mixed and equally accessible, and where there is a high degree of spatial rigidity within cell populations. It is also easy to alter other assumptions underlying our model. For instance, our conclusions regarding acetyl-CoA being the driving resource depend on an assumption we made in building the model that G cells consume the resource. While this is biologically plausible, mathematically we could have assumed the opposite, namely that the resource is consumed by cells in the Q state and not by cells in the G state. In that case too our model could give similar oscillations - switching rates would still need to be density/resource dependent, but the form of dependence on resource would need to be reversed so that the high and low-q branches would be the mirror images, with the low *q* branch being the only one at low *a* and the high *q* branch being the only one at large values of *a*. And hence the waveform of the oscillating resource would be flipped compared to Figures 2 and 3 - i.e., when *q* is high *a* would be decreasing, while when *q* is low *a* would be increasing. If one could find a metabolite that exhibited this waveform, then that metabolite would be an equally likely possibility as a driver of the Q-G transition, except that it would have to act such that it caused a switch from Q to G when it crossed a low threshold, or caused the opposite transition when it crossed a high threshold. From the considerable data available, we have not found a reasonable molecule with such a reversed waveform. Moreover, we know of no process which consumes a metabolite in the Q state in the way described, so for now acetyl-CoA driving the Q-G transition and being consumed during growth is the most parsimonious explanation. Nevertheless, this shows how our framework could be easily used in alternate scenarios.

Currently, existing experimental approaches to study such metabolically-driven Q-G oscillations are very limited. Crude readouts, such as oxygen consumption, have very limited resolution even to show the Q and G states, as the bistability begins to break down. Gene expression analysis (even when done in single cells) is a late, end-point readout which cannot explain this bistability but instead occurs after a switch. The key to experimentally studying such bistability, therefore, will be the development of in vivo intracellular metabolic sensors with excellent dynamic range and sensitivity, for metabolites like acetyl-CoA or NADPH. This will allow the development of more precise models to predict commitment steps, and identify differences within the population of cells, that will help understand reversibility (between states), hysteresis and other apparent phenomena.

## Methods

### Experimental methods and data sets

Chemostat culture and cell division datasets: All dO2 data were obtained from YMCs set up similar to already published data (Tu *et al.*, 2005, 2007; Kudlicki *et al.*, 2007; Mohler *et al.*, 2008). In these studies, yeast cells were grown in chemostat cultures using semi-defined medium, and yeast metabolic cycles were set up as described earlier (Tu *et al.*, 2005; Tu, 2010). Data for cell division across three metabolic cycles was obtained from earlier studies (Tu *et al.*, 2005; Laxman *et al.*, 2010). YMC gene expression and metabolite datasets: Gene expression datasets were obtained from (Tu *et al.*, 2005; Kudlicki *et al.*, 2007), and metabolite oscillation datasets were obtained from (Tu *et al.*, 2007; Mohler *et al.*, 2008; Cai *et al.*, 2011; Machné and Murray, 2012), including acetyl-CoA oscillation datasets.

### Parameter values and their q/a dependencies

Figures 2C, 3C(i), 4A(ii), 4B(ii), 5A and 5B (default choices):

To produce the oscillation shown in these figures, we make the following choices (within scenario 3c):

γ=1.665 hr^−1^. σ=0.3996 hr^−1^, μ=1, ν_GQ_=16.65 hr^−1^, ν_QG_=*h(q)*, where *h(q)* is the Hill function *h(q)* = ν_m_(1+β(*q/K*)^20^)/(1+(*q/K*)^20^) with β=0.01, *K*=*a*^2^/(0.75^2^+*a*^2^), ν_m_=16.65×(1.65-1.25K). We use this Hill function with such a high Hill coefficient to approximate a step function which drops rapidly from ν_m_ to βν_m_ at *q*=*K*.

Figure 4, other panels:

The other panels of Fig 4 are made using exactly the same equations and parameter choices as above, except for varying *σ* and *γ* as mentioned in the Fig 4 caption.

Figure 3C, other panels:

As above, except that

i. ½_m_=16.65×(1.65-1.25K)×2.25K hr^−1^ and σ=0.3596 hr^−1^.
ii. ½_m_=16.65×(1.65-1.25K)+16.651.85*a*^10^/(200+*a*^10^) hr^−1^ and σ=0.3297 hr^−1^.
iii. ½_m_=16.65×2.25K’ hr^−1^ and σ=0.3397 hr^−1^. *σ* values were varied in order to keep the time period close to 4 hours.

Figure 5C, addition of bolus:

Until time *t* = 11.5 hours, the simulation is the same as in Fig 2C. At *t* = 11.5 hrs, the resource level is abruptly changed to 1.75 (just above its peak value in previous cycles, which was 1.73), and then the simulation is continued with the same equations and parameter values.

In all the above cases, the simulations were started, at *t* = 0 hours, with initial conditions q=1 and *a*=10^−6^ (i.e., we start with all cells in a quiescent state and starved of the resource). Simulations and figures were produced in Matlab. We used the ode45 differential equation integrator. The code is provided in Supplemental material. As extra controls, we checked that the stiff solver ode15s also provided the same results for the simulations in Fig 2 and 5, and a Mathematica notebook which repeats many of the same simulations, using the default NDSolve algorithm within Mathematica, is also provided with the Supplemental material.

## Acknowledgements

SK is supported by funds from the Simons Foundation, and institutional support from NCBS-TIFR. SL is supported by a Wellcome Trust-DBT IA Intermediate Fellowship (IA/I/14/2/501523), and institutional support from inStem and the Department of Biotechnology.

